# An infection and pathogenesis mouse model of SARS-CoV-2-related pangolin coronavirus GX_P2V(short_3UTR)

**DOI:** 10.1101/2024.01.03.574008

**Authors:** Lai Wei, Shuiqing Liu, Shanshan Lu, Shengdong Luo, Xiaoping An, Huahao Fan, Weiwei Chen, Erguang Li, Yigang Tong, Lihua Song

## Abstract

SARS-CoV-2-related pangolin coronavirus GX_P2V(short_3UTR) is highly attenuated, but can cause mortality in a specifically designed human ACE2-transgenic mouse model, making it an invaluable surrogate model for evaluating the efficacy of drugs and vaccines against SARS-CoV-2.

## INTRODUCTION

Two SARS-CoV-2-related pangolin coronaviruses, GD/2019 and GX/2017, were identified prior to the COVID-19 outbreak (1, 2). The respective isolates, termed pCoV-GD01 and GX_P2V, were cultured in 2020 and 2017, respectively (2, 3). The infectivity and pathogenicity of these isolates have been studied (4-6). The pCoV-GD01 isolate, which has higher homology with SARS-CoV-2, can infect and cause disease in both golden hamsters and hACE2 mice (4). In contrast, GX_P2V was found to be highly attenuated in previously tested animals like golden hamsters, BALB/c mice and two types of human ACE2-transgenic (hACE2) mouse (5, 6). We previously reported that the early passaged GX_P2V isolate was actually a cell culture-adapted mutant, named GX_P2V(short_3UTR), which possesses a 104-nucleotide deletion at the 3’-UTR (6). In this study, we analyzed its adaptive mutation in cell culture, and assessed its pathogenicity in a unique CAG-hACE2 mouse model. We found that GX_P2V(short_3UTR) can infect hACE2 mice, with high viral loads detected in both lung and brain tissues, which are correlated with the strong expression of hACE2 in these tissues. This infection resulted in 100% mortality in the hACE2 mice. We surmise that the cause of death may be linked to the occurrence of late brain infection.

## RESULTS

We first analyzed the adaptive mutations of the GX_P2V(short_3UTR) mutant in cell cultures by random cloning and sequencing. The passaged mutant was cloned through two successive plaque assays. Eight viral clones were chosen for next-generation sequencing (National Genomics Data Center of China, GSA: CRA014225). These clones, when compared with the genome of the original mutant (6), all shared four identical mutations: ORF1ab_D6889G, S_T730I, S_K807N, and E_A22D (Supporting Information, Table S1). Clone 7, named as GX_P2V C7, was randomly selected for the evaluation of viral pathogenicity in hACE2 mice (Figure 1A). The hACE2 mouse model expressing human ACE2 under control of the CAG promoter was developed using random integration technology by Beijing SpePharm Biotechnology Company.

**Figure 1:**
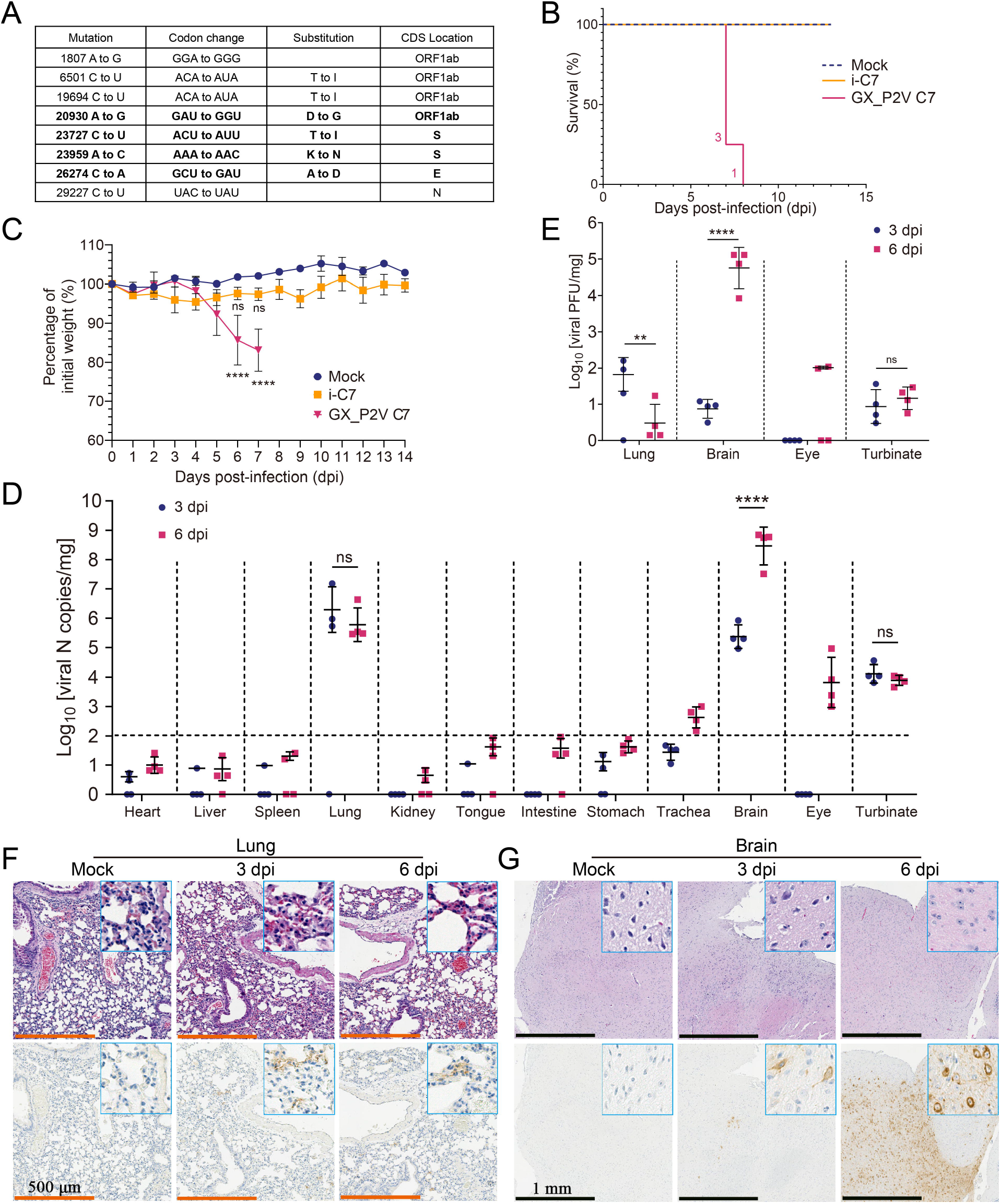
Characterization of a lethal infection model in human ACE2-transgenic mice caused by the attenuated SARS-CoV-2-related pangolin coronavirus GX_P2V C7. **A** Mutations in GX_P2V C7 compared to the GX_P2V(short_3UTR) isolate (NCBI accession number: MW532698). The four identical mutations are shown in bold. **B** Survival of hACE2 transgenic mice following intranasal infection with GX_P2V C7 (*n* = 4), inactivated GX_P2V C7 (i-C7, *n* = 4), and mock infection (*n* = 4). The number of deceased mice on each specific day is annotated on the left of the survival curve. **C** Percentage of initial weight of hACE2 transgenic mice after intranasal infection with GX_P2V C7 (*n* = 4), i-C7 (*n* = 4), and mock infection (*n* = 4). The statistical significance of the differences between mock-infected (*n* = 4, blue dots) and GX_P2V C7-infected (*n* = 4, red dots) or i-C7-infected mice (*n* = 4, orange dots) at 6 or 7 dpi are shown. The error bars represent the means ± SDs. **D** Quantification of GX_P2V N gene copies in heart, liver, spleen, lung, kidney, tongue, intestine, stomach, trachea, brain, eye, and turbinate homogenates at 3- and 6-day post-infection (dpi) (*n* = 4 per group). The limit of detection (LOD) for viral RNA loads in the original samples was Log_10_[10^2^ copies/mg]. The error bars represent the means of Log_10_[copies/mg] ± SDs. The significances of the comparisons in the lung, brain, and turbinate are shown. **E** Infectious viral titers in lung, brain, eye, and turbinate homogenates were measured by plaque forming assay at 3 and 6 dpi (*n* = 4 per group). The statistical significance of the differences in the lung, brain, and turbinate are shown. The error bars represent means of Log_10_[pfu/mL] ± SDs. **F, G** Hematoxylin and eosin (H&E) staining and immunohistochemical (IHC) staining with an anti–SARS-CoV-2 N-specific antibody (SARS-CoV-2) revealed viral antigen–positive cells (brown) in the lung (**F**) and brain (**G**), as shown at high magnification in the inset. Scale bars, 500[:μm (**F**) and 1 mm (**G**), respectively. ^*^*P* < 0.05, ^**^*P* < 0.01, ^***^*P* < 0.001, ^****^*P* < 0.0001, *P* > 0.05, not significant (ns); two-way ANOVA followed by Sidak’s multiple comparison test.

Next, we assessed whether GX_P2V C7 could cause disease in hACE2 mice by monitoring daily weight and clinical symptoms. A total of four 6 to 8-week-old hACE2 mice were intranasally infected with a dosage of 5×10^5^ plaque-forming units (pfu) of the virus. Four mice inoculated with inactivated virus and four mock-infected mice were used as controls. Surprisingly, all the mice that were infected with the live virus succumbed to the infection within 7-8 days post-inoculation, rendering a mortality rate of 100% (Figure 1B). The mice began to exhibit a decrease in body weight starting from day 5 post-infection, reaching a 10% decrease from the initial weight by day 6 (Figure 1C). By the seventh day following infection, the mice displayed symptoms such as piloerection, hunched posture, and sluggish movements, and their eyes turned white. The criteria for clinical scoring of the mice and the daily clinical scores post-infection with GX_P2V C7 are provided in the Supporting Information, Figure S1.

We then evaluated the tissue tropism of GX_P2V C7 in hACE2 mice. Using the infection method described above, eight hACE2 mice were infected, eight mice were inoculated with inactivated virus, and another eight mock-infected mice were used as controls. The organs of four randomly selected mice in each group were dissected on days 3 and 6 post-infection for quantitative analysis of viral RNA and titer. We detected significant amounts of viral RNA in the brain, lung, turbinate, eye, and trachea of the GX_P2V C7 infected mice (Figure 1D), whereas no or a low amount of viral RNA was detected in other organs such as the heart, liver, spleen, kidneys, tongue, stomach, and intestines. Specifically, in lung samples, we detected high viral RNA loads on days 3 and 6 post-infection, with no significant difference between these two time points (∼ 6.3 versus ∼ 5.8 Log_10_[copies/mg]). In brain samples, on day 3 post-infection, viral RNA was detected in all four infected mice, with an average value of 5.4 Log_10_[copies/mg]. Notably, by day 6 post-infection, we detected exceptionally high viral RNA loads (∼ 8.5 Log_10_[copies/mg]) in the brain samples from all four infected mice (Figure 1D). On days 3 and 6 post-infection, the viral RNA loads in the turbinate were similar, approximately 4.1 and 3.9 Log_10_[copies/mg], respectively. The viral RNA loads in the trachea and eyes of the mice surpassed the limit of detection only on day 6 post-infection, with values of 2.6 and 3.8 Log_10_[copies/mg], respectively. Regarding the infectious viral titers, lung tissues at day 3 post-infection had a value of ∼ 1.8 Log_10_[pfu/mg], which decreased to ∼ 0.5 Log_10_[pfu/mg] by day 6. Importantly, the highest infectious titers were detected in the brain on day 6, which was significantly greater than that on day 3 (∼ 0.9 vs ∼ 4.8 Log_10_[pfu/mg]) (Figure 1E). Additionally, there were no significant differences in the infectious titers in the turbinate between day 3 (∼ 0.9 Log_10_[pfu/mg]) and day 6 (∼ 1.2 Log_10_[pfu/mg]) (Figure 1E). By day 6, approximately 2.0 Log_10_[pfu/mg] was detected in the eyes of two mice. Neither inactivated GX_P2V C7 nor mock infection caused death or any clinical symptoms in the mice (Figure 1B-C and Supporting Information, Figure S2). In summary, in the mice infected with live virus, the viral load in the lungs significantly decreased by day 6; both the viral RNA loads and viral titers in the brain samples were relatively low on day 3, but substantially increased by day 6. This finding suggested that severe brain infection during the later stages of infection may be the key cause of death in these mice.

To determine the mechanisms underlying GX_P2V C7-induced death in hACE2 mice, we examined the pathological changes, presence of viral antigens, and cytokine profiles in the lung and brain tissues of the mice on days 3 and 6 post-infection (Figure 1F-G, and Supporting Information, Figure S3 and S4). On both days, compared to those of control mice, the lungs of infected mice showed no significant pathological alterations, with only minor inflammatory responses due to slight granulocyte infiltration (Figure 1F). On day 3 post-infection, shrunken neurons were visible in the cerebral cortex of the mice. By day 6, in addition to the shrunken neurons, there was focal lymphocytic infiltration around the blood vessels, although no conspicuous inflammatory reaction was observed (Figure 1G). Upon staining for viral nucleocapsid protein via immunohistochemistry, viral antigens were detected in both the lungs and brains on days 3 and 6 post-infection, with extensive viral antigens notably present in the brain on day 6 (Figure 1F-G). These findings align with the viral RNA load assessments in the lung and brain tissues (Figure 1D). We also performed a Luminex cytokine assay to detect 31 cytokines/chemokines in the lung and brain tissues of the mice (Supporting Information, Figure S3 and S4). Consistent with the pathological findings, there were slight increases or decreases in the levels of many cytokines/chemokines in lung and brain tissues compared to those in control tissues, but the levels of key inflammatory factors, such as IFN-γ, IL-6, IL-1β, and TNF-α, did not significantly change. In brief, these analyses revealed that GX_P2V C7 infection in hACE2 mice did not lead to severe inflammatory reactions, a finding that aligns with previous reports by Zhengli Shi’s group using GX_P2V infection in two different hACE2 mouse models (5), as well as our own findings in the golden hamster model (6).

## DISCUSSION

To the best of our knowledge, this is the first report to analyze the cell-adapted mutations of pangolin coronavirus GX_P2V, and to show it can cause mortality in hACE2 mice. Our findings are evidently inconsistent with those of Zhengli Shi *et al*. (5), who tested the virulence of GX_P2V in two different hACE2 mouse models. It is very likely that the high pathogenicity of GX_P2V C7 in our hACE2 mice is due to the strong expression of hACE2 in the mouse brain. Under normal circumstances, both human and mouse brains exhibit low expression of ACE2 (6, 7). Furthermore, while the company has not yet published a paper detailing the construction and characterization of this hACE2 mouse model, we are notified that these hACE2 mice have abnormal physiology, as indicated by relatively reduced serum triglyceride, cholesterol, and lipase levels, compared to those of wild-type C57BL/6J mice. Thus, the outcomes from the mouse infections in this study have no correlation with human infections, and do not alter the fundamental nature of GX_P2V(short_3UTR) as being highly attenuated.

Currently, there is an urgent need for the development of broadly protective vaccines against pan-SARS-CoV-2, yet the emergence of the next SARS-CoV-2 variant is unpredictable. The pangolin coronavirus GX_P2V(short_3UTR), which shares a certain degree of homology with SARS-CoV-2, may be valuable in assessing the effectiveness of broad-spectrum COVID-19 vaccine candidates against unknown future variants. Moreover, our lethal mouse infection model presents no obvious inflammatory responses in the main affected organs, the lungs and brain, thereby providing an alternative model to evaluate antiviral drugs’ efficacy in inhibiting viral replication *in vivo*. In summary, our study provides a unique perspective on the pathogenicity of GX_P2V and offers an invaluable model for assessing the efficacy of drugs and vaccines against SARS-CoV-2.

## ACKNOWLEDGEMENTS

This work was supported by NSFC-MFST project (China–Mongolia) (grant number 32161143027), National Key R&D Program of China (2021YFC2301804) and Biosafety Special Program (No. 19SWAQ 13).

## ETHICS STATEMENT

All animals involved in this study were housed and cared for in an AAALAC (Association for Assessment and Accreditation of Laboratory Animal Care) accredited facilities. The procedure for animal experiments (IACUC-2019-0027) was approved by the Institutional Animal Care and Use Committee of the Fifth Medical Center, General Hospital of the Chinese People’s Liberation Army, and complied with IACUC standards.

## AUTHOR CONTRIBUTIONS

L.Song conceived and designed the study and wrote the manuscript. L.W., S.Liu, S.Lu., and S.Luo. performed the experiments and analyzed the data. X.A., H.F., W.C., E.L. and Y.T. analyzed the data and edited the manuscript. L.W. and L.Song wrote the manuscript and all the authors approved the manuscript.

## CONFLICT OF INTERESTS

The authors declare no competing interests.

## SUPPORTING INFORMATION

Additional Supporting Information for this article can be found online at

## DATA AVAILABILITY

All the data supporting the findings of this study are available within the article and the Supporting Information, or from the corresponding author upon reasonable request.

## Notes

### Competing Interest Statement

The authors have declared no competing interest.

### Summary of Updates

The title, abstract and discussion parts were fundamentally updated. The previous manuscript may mislead readers to believe that the attenuated pangolin coronavirus GX_P2V(short_3UTR) posed a spillover risk to human brains, resulting in a 100% mortality rate, which sparked panic among the public. In this revision, we revised the manuscript to state the fact that these animal outcomes cannot be applicable to humans, but this infection model is invaluable for evaluating the efficacy of drugs and vaccines against SARS-CoV-2.

