## Supporting Information for "An infection and pathogenesis mouse model of SARS-CoV-2-related pangolin coronavirus GX_P2V(short_3UTR)"

### MATERIALS AND METHODS

#### Cell lines and viruses

The Vero and BGMK cell lines were obtained from the American Type Culture Collection (ATCC) and grown in minimum essential medium (MEM) (HyClone, USA) supplemented with 10% (vol/vol) fetal bovine serum (FBS) (PAN, USA) and 1% (vol/vol) penicillin/streptomycin (100 IU/mL and 100 mg/mL, respectively) (Gibco, USA) at 37°C with 5% $CO_2$. The pangolin coronavirus GX_P2V(short_3UTR), originally cultured from the lung-intestine mixed samples of a pangolin captured in anti-smuggling operations in 2017, was passaged in Vero cells (1). Briefly, Vero cells were infected with GX_P2V(short_3UTR) at a multiplicity of infection (MOI) of 0.01 for 2 hours, followed by a double rinse with phosphate-buffered saline (PBS). Then, the Vero cells were cultured in MEM supplemented with 2% FBS and 1% penicillin/streptomycin (Gibco, USA) for 48 hours. Subsequently, the supernatant was harvested, aliquoted, and stored at -80°C.

#### Plaque cloning of GX_P2V(short_3UTR)

BGMK cells from a T175 flask were seeded at a ratio of 1:4 in six-well plates. The following day, the medium was discarded, and the cells were infected with 20 plaque-forming units (PFUs) per well of GX_P2V(short_3UTR), suspended in 500 μL of MEM supplemented with 2% FBS. At 2 hours post-infection (hpi) at room temperature, the viral inoculum was removed, and each well was filled with 3 mL of 1% (wt/vol) methylcellulose overlay (2×MEM and 2% methylcellulose mixed at a ratio of 1:1). The plates were then incubated at 37°C in a 5% $CO_2$ incubator. Following a five-day incubation period, the plates were carefully handled to avoid disturbing the cell overlay, and the locations of viral plaques were identified and marked with a marker under a light source. Using a flat-head pipette tip, we meticulously reached each marked plaque site. The tip was then inserted to the bottom of the plate to aspirate approximately 20 μL of the semi-solid medium containing the viral plaques. This aspirated medium was then transferred into 1 mL of MEM. The mixture was thoroughly vortexed to ensure uniform distribution of the virus particles. Subsequently, Vero cells, seeded in a 24-well plate, were infected with this viral suspension to facilitate the expansion of the virus. Following this initial round of cloning, a successive plaque assay was performed to further clone the virus. Ultimately, a total of eight clones were isolated through two successive rounds of plaque assays.

#### Next-generation sequencing

Next-generation sequencing (NGS) was used to analyze a total of eight GX_P2V(short_3UTR) clones. The viral RNAs were extracted using an SE Viral DNA/RNA Kit (Omega, USA). The sequencing libraries were constructed with the NEBNext® Ultra™ II Directional RNA Library Prep Kit for Illumina® (NEB, USA). The NGS service used was provided by Annoroad, a commercial company based in Beijing. The obtained sequence data were mapped to the reference sequence of GX_P2V(short_3UTR) (NCBI accession number: MW532698) and mutations were identified using Geneious Prime software. The raw sequence data of these eight viral clones have been deposited in the Genome Sequence Archive in National Genomics Data Center, China National Center for Bioinformation (GSA: CRA014225) that are publicly accessible at https://ngdc.cncb.ac.cn/gsa.

#### Mouse infection experiments

Six-to-eight-week-old C57BL/6J CAG-hACE2 mice under specific pathogen-free (SPF) conditions were purchased from SpePharm Biotechnology (Beijing, China). The mice were housed in individually ventilated cages (IVCs) and fed standard chow. After deep anesthesia was induced by intraperitoneal injection of pentobarbital (50 mg of pentobarbital/kg of mouse body weight), the mice were intranasally infected with $5\times10^5$ PFU of infectious GX_P2V C7, $5\times10^5$ PFU of heat inactivated GX_P2V C7 (i-C7) or 2% FBS MEM (mock) in a 20 μL volume. Initially, the daily weight and clinical symptoms of the GX_P2V C7-infected group (n=4), the i-C7-infected group (n=4), and the mock-infected group (n=4) were monitored. Then, the mice in the GX_P2V C7-infected group (n=4), the i-C7-infected group (n=4), and the mock-infected group (n=4) were euthanized on the third day post-infection (3 dpi), and an additional series of three groups were euthanized at 6 dpi.

Following euthanasia, the left brain and left lung of the mice were rapidly fixed in 4% paraformaldehyde (Solarbio, China) for histopathological analysis and immunohistochemistry. The remaining tissues (heart, liver, spleen, right lung, kidney, tongue, intestine, stomach, right brain, and trachea) were weighed and then submerged in 800 μL of sterile PBS and homogenized in a cryomiller. After centrifugation at 12,000 rpm for 10 min, the supernatant was aliquoted and stored at -80°C for analysis.

**Infectious titer determination by plaque assay**

BGMK cells from a T175 flask were initially seeded in six-well plates at a ratio of 1:4. On the subsequent day, the medium in each well was replaced with 500 μL of serially diluted tissue homogenates. These homogenates were prepared in MEM supplemented with 2% FBS. At 2 hpi at room temperature, 3 mL of 1% (wt/vol) methylcellulose overlay was added to each well. The plates were then incubated for a duration of five days at 37°C in a 5% $CO_2$ incubator. Post-incubation, the plates underwent fixation using 4% polyformaldehyde (Solarbio, China) for 20 minutes. Following fixation, the plates were stained with 0.1% (wt/vol) crystal violet (Solarbio, China). After staining, the plates were thoroughly rinsed with deionized water to remove excess dye and to enhance the clarity of the plaques. Finally, the plaques were counted to determine the infectious titer of the virus present in the tissue homogenates.

**RT–qPCR analysis**

Viral RNA was extracted from the supernatant of 200 μL of tissue homogenates using a cell/tissue total RNA extraction kit (Nobelab, China). Total RNA was reverse transcribed with a HiScript III RT SuperMix for qPCR (+gDNA wiper) kit (Vazyme, China). Reverse transcription was performed at 37°C for 15 min, followed by 85°C for 5 sec. QuantiNova PCR kits (Qiagen, Germany) for quantifying the N gene copy numbers were used for 40 cycles (15 sec at 95°C and 1 min at 60°C), with the following primers: TCTTCCTGCTGCAGATTTGGAT, reverse primer: TTACACATTAGGGCTCTTCCATATAGG and probe (FAM-TGCAGACCACACAAGGCAGATGGGC-TAMRA). Primers and probes were used at final concentrations of 200 nM and 100 nM, respectively. A standard plasmid of the targeted fragments was used for quantitative analysis, and the limit of detection (LOD) was set at 40 copies per reaction.

**Chemokine and cytokine protein assays**

Lung and brain homogenates were incubated with Triton X-100 (at a 1% final concentration) for 1 hour at room temperature to inactivate the infectious viral particles. Cytokine and chemokine protein levels were measured using the Bioplex 200 system (Bio-Rad, USA) platform with the Bio-Plex Pro$^{TM}$ Mouse Chemokine Assay (Cat. 10000057971, Bio-Rad, USA) following the manufacturer's instructions.

### Histology and immunohistochemistry

The left hemisphere and left lung were fixed in 4% paraformaldehyde and embedded in paraffin. Paraffin sections (approximately 4 μm thick) were stained with hematoxylin-eosin (H&E) as described previously (1). Histopathological results were analyzed by professional pathologists.

To detect the GX_P2V C7 virus antigen, paraffin sections were deparaffinized in xylene and rehydrated in a graded ethanol series. To restore antigenicity, the slides were submerged in ethylenediaminetetraacetic acid (EDTA) antigen repair solution (Cat. ZLI-9072, ZSGB, China) in a microwaveable vessel. Then, the vessel was placed inside the microwave and boiled for 20 minutes. After that, the vessel was removed and washed with cold PBS for 10 minutes. Endogenous peroxidase was blocked using 3% hydrogen peroxide (Cat. PV-6000D, ZSGB, China) for 20 minutes at room temperature (RT). The slides were washed in PBS, and nonspecific binding was blocked by incubating the slides in 10% normal goat serum for 30 minutes at RT. Then, the sections were incubated overnight at 4°C (1:1000 dilution in 1% BSA in PBS) with a mouse anti-SARS-CoV-2 N protein monoclonal antibody (HENDERSON, China). The secondary antibody used was an HRP-labeled goat anti-mouse/rabbit IgG (the Mouse/Rabbit Polymer Method Detection System kit, Cat. PV-6000D, ZSGB, China), which was incubated with the slides at 37°C in the dark for 30 minutes. The diaminobenzidine tetrahydrochloride (DAB) substrate (Cat. PV-6000D, ZSGB, China) was used for visualization. Before the organ tissues were analyzed via microscopy (Nikon, Japan), the sections were counterstained with hematoxylin, dehydrated in a graded ethanol series, cleared in xylene, and mounted with permanent mounting medium and coverslips.

**Supporting Information Table S1. Mutations in eight GX_P2V(short_3UTR) clones compared to the parent GX_P2V(short_3UTR) (NCBI accession number: MW532698).**

| Name | Mutation | Codon Change | Substution |
|---|---|---|---|
| GX_P2V(short_3UTR) Clone1 | A20930G | GAU -> GGU | D > G |
| | C20932U | CUU -> UUU | L > F |
| | C23727U | ACU -> AUU | T > I |
| | A23959C | AAA -> AAC | K > N |
| | C26274A | GCU -> GAU | A > D |
| | C29227U | UAC -> UAU | |
| GX_P2V(short_3UTR) Clone2 | C14290G | CAA -> GAA | Q > E |
| | C18905U | CCU -> CUU | P > L |
| | A20930G | GAU -> GGU | D > G |
| | U23006A | UAU -> AAU | Y > N |
| | C23727U | ACU -> AUU | T > I |
| | A23959C | AAA -> AAC | K > N |
| | C26274A | GCU -> GAU | A > D |
| | A29323U | CCA -> CCU | |
| GX_P2V(short_3UTR) Clone3 | U12103C | UAU -> UAC | |
| | A20930G | GAU -> GGU | D > G |
| | C23727U | ACU -> AUU | T > I |
| | A23959C | AAA -> AAC | K > N |
| | C26274A | GCU -> GAU | A > D |
| GX_P2V(short_3UTR) Clone4 | A20930G | GAU -> GGU | D > G |
| | U21390C | CUU -> CUC | |
| | C23727U | ACU -> AUU | T > I |
| | A23959C | AAA -> AAC | K > N |
| | C26274A | GCU -> GAU | A > D |
| GX_P2V(short_3UTR) Clone5 | C337U | CGC -> CGU | |
| | G9055U | GUG -> GUU | |
| | A20930G | GAU -> GGU | D > G |
| | C23727U | ACU -> AUU | T > I |
| | C23797U | GGC -> GGU | |
| | A23959C | AAA -> AAC | K > N |
| | C26274A | GCU -> GAU | A > D |
| | C29251U | GAC -> GAU | |
| GX_P2V(short_3UTR) Clone6 | U4106C | GUU -> GUC | |
| | G15076A | GCU -> ACU | A > T |
| | U15282C | UGU -> UGC | |
| | A20930G | GAU -> GGU | D > G |
| | A23035U | CCA -> CCU | |

| | | | |
|---|---|---|---|
| | C23727U | ACU -> AUU | T > I |
| | A23959C | AAA -> AAC | K > N |
| | C26274A | GCU -> GAU | A > D |
| GX_P2V(short_3UTR) Clone7 | A1807G | GGA -> GGG | |
| | C6501U | ACA -> AUA | T > I |
| | C19694U | ACA -> AUA | T > I |
| | A20930G | GAU -> GGU | D > G |
| | C23727U | ACU -> AUU | T > I |
| | A23959C | AAA -> AAC | K > N |
| | C26274A | GCU -> GAU | A > D |
| | C29227U | UAC -> UAU | |
| GX_P2V(short_3UTR) Clone8 | A1807G | GGA -> GGG | |
| | C6501U | ACA -> AUA | T -> I |
| | C19694U | ACA -> AUA | T -> I |
| | A20930G | GAU -> GGU | D -> G |
| | C23727U | ACU -> AUU | T -> I |
| | A23959C | AAA -> AAC | K -> N |
| | C26274A | GCU -> GAU | A -> D |
| | C29227U | UAC -> UAU | |

**Supporting Information Figure S1**

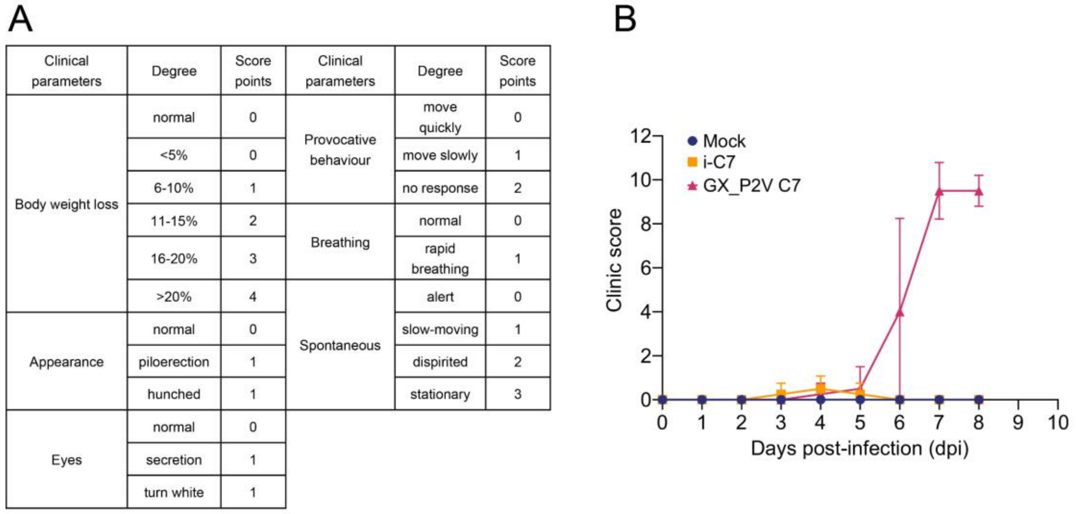

A

| Clinical parameters | Degree | Score points | Clinical parameters | Degree | Score points |
|---|---|---|---|---|---|
| Body weight loss | normal | 0 | Provocative behaviour | move quickly | 0 |
| | <5% | 0 | | move slowly | 1 |
| | 6-10% | 1 | | no response | 2 |
| | 11-15% | 2 | Breathing | normal | 0 |
| | 16-20% | 3 | | rapid breathing | 1 |
| | >20% | 4 | Spontaneous | alert | 0 |
| Appearance | normal | 0 | | slow-moving | 1 |
| | piloerection | 1 | | dispirited | 2 |
| | hunched | 1 | | stationary | 3 |
| Eyes | normal | 0 | | | |
| | secretion | 1 | | | |
| | turn white | 1 | | | |



**Supporting Information Figure S1. Clinical scores of GX_P2V C7-infected, inactivated GX_P2V C7-infected and mock-infected CAG-hACE2 transgenic mice (n=4 per group) (B) and the standard of scoring (A).** Inactivated GX_P2V C7 (i-C7). The error bars represent the means ± SDs.

**Supporting Information Figure S2**

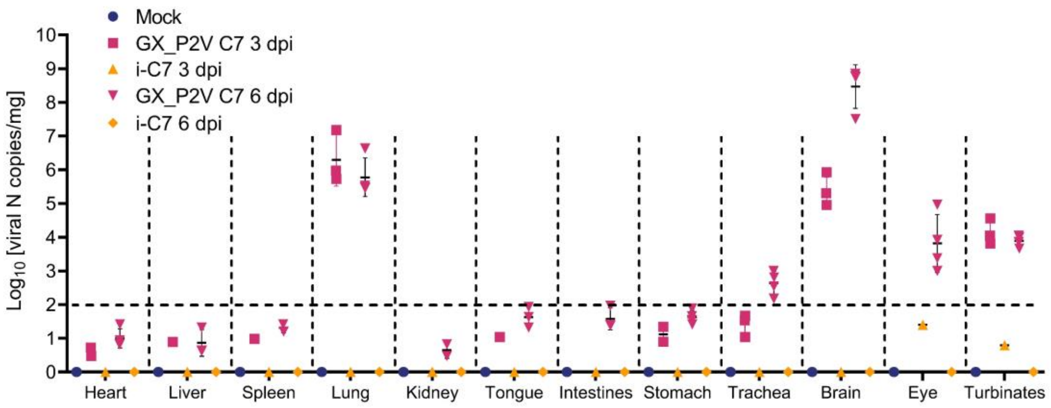


**Supporting Information Figure S2. Viral loads in different tissue homogenates on days 3 and 6 post infection (3 and 6 dpi) (n=4 per group).** Inactivated GX_P2V C7 (i-C7). The error bars represent the means of $Log_{10}$[copies/mg] ± SDs.

**Supporting Information Figure S3**

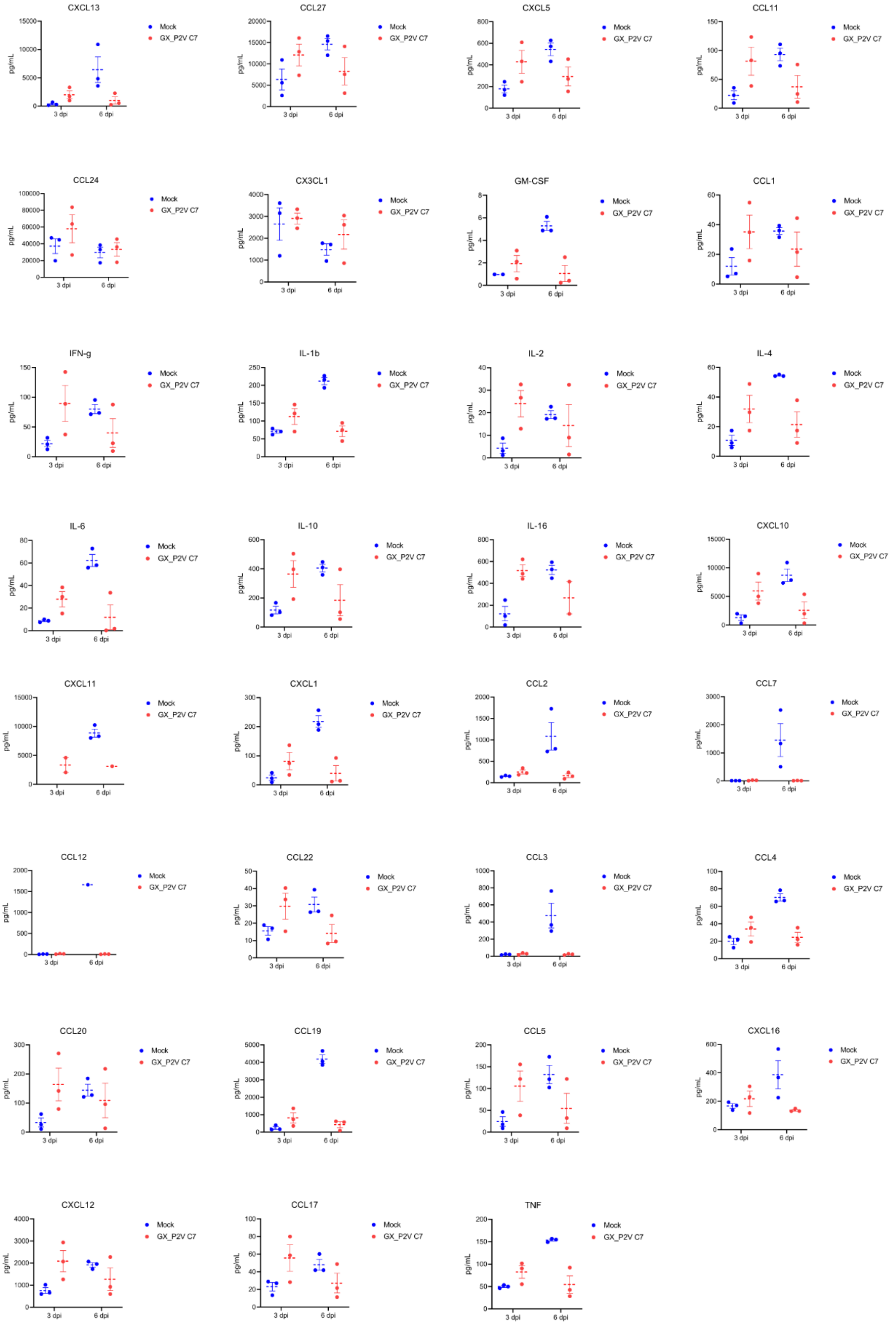


**Supporting Information Figure S3. Cytokine and chemokine protein levels in the brain tissues of GX_P2V C7 and mock-infected CAG-hACE2 mice at 3 and 6 dpi (n=3 per group) were measured via a multiplex platform.** The error bars represent the means ± SDs.

**Supporting Information Figure S4**

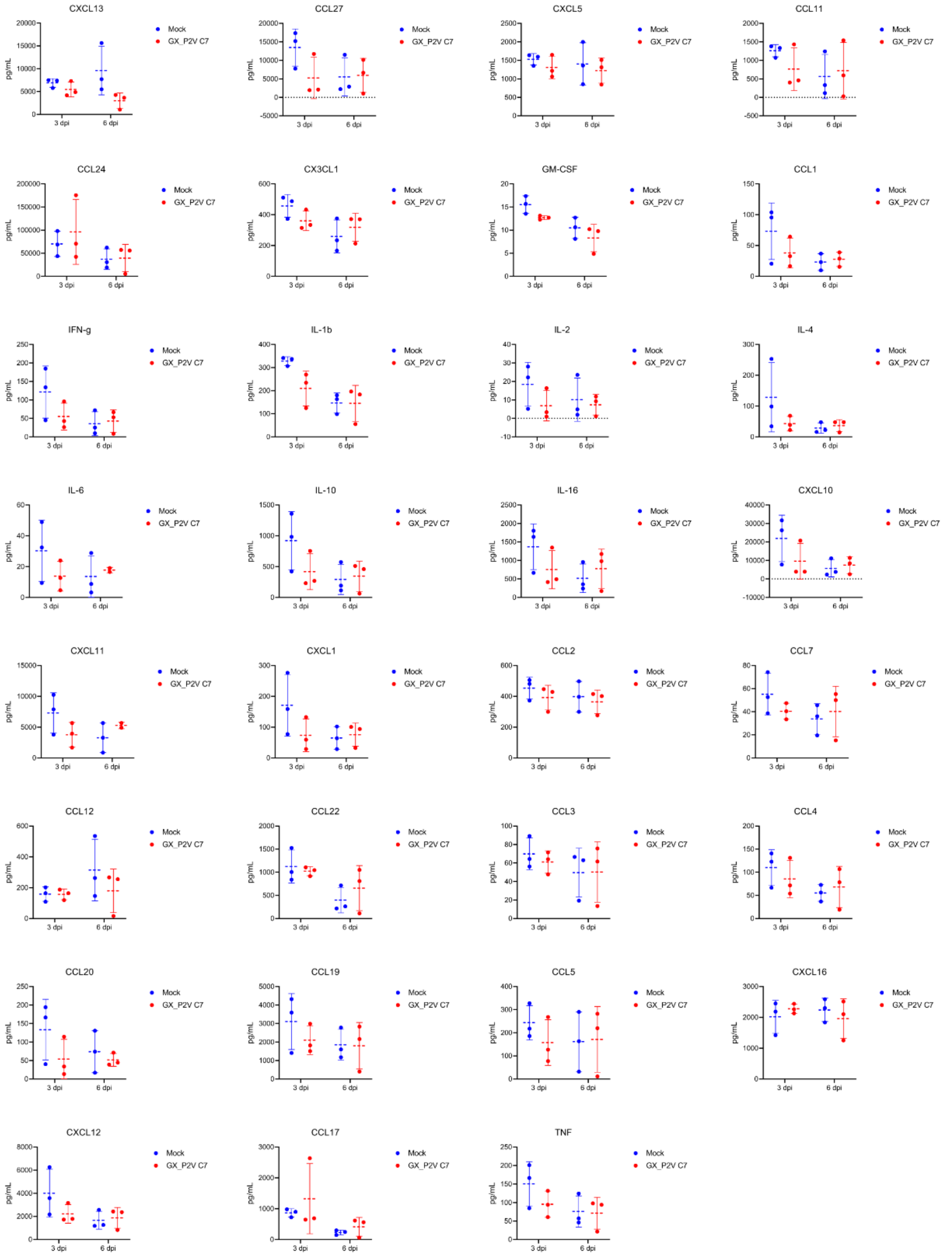


**Supporting Information Figure S4. Cytokine and chemokine protein levels in lung tissue from GX_P2V C7 and mock-infected CAG-hACE2 mice at 3 and 6 dpi (n=3 per group) were measured via a multiplex platform.** The error bars represent the means ± SDs.
